# Comprehensive single cell RNAseq analysis of the kidney reveals novel cell types and unexpected cell plasticity

**DOI:** 10.1101/203125

**Authors:** Jihwan Park, Rojesh Shrestha, Chengxiang Qiu, Ayano Kondo, Shizheng Huang, Max Werth, Mingyao Li, Jonathan Barasch, Katalin Suszták

## Abstract

A key limitations to understand kidney function and disease development has been that specific cell types responsible for specific homeostatic kidney function or disease phenotypes have not been defined at the molecular level.

To fill this gap, we characterized 57,979 cells from healthy mouse kidneys using unbiased single-cell RNA sequencing. We show that genetic mutations that present with similar phenotypes mostly affect genes that are expressed in a single unique differentiated cell type. On the other hand, we found unexpected cell plasticity of epithelial cells in the final segment of the kidney (collecting duct) that is responsible for final composition of the urine. Using computational cell trajectory analysis and *in vivo* linage tracing, we found that, intercalated cells (that secrete protons) and principal cells (that maintain salt, water and potassium balance) undergo a Notch mediated interconversion via a newly identified transitional cell type. In disease states this transition is shifted towards the principal cell fate. Loss of intercalated cells likely contributes to metabolic acidosis observed in kidney disease.

In summary, single cell analysis advanced a mechanistic description of kidney diseases by identifying a defective homeostatic cell lineage.

**One Sentence Summary:** A comprehensive single cell atlas of the kidney reveals a transitional cell type and cell plasticity determined by Notch signaling which is defective in chronic kidney disease.

## Main Text

The kidney filters the blood to remove nitrogen, water and other waste products. Furthermore, the kidneys control electrolyte concentrations, acid-base balance, secrete hormones and regulate blood pressure. The glomerulus produces the primary filtrate from the plasma. Majority of water and electrolytes undergo an unregulated reabsorption in the renal proximal tubules, while some substrates are still secreted into the filtrate. The next segment (Loop of Henle) is mostly involved solute concentration. The distal tubule and finally the collecting duct are the kidney segments where regulated solute transport occurs and therefore these segments are critical for maintaining electrolyte and water homeostasis. Kidney cells have previously been annotated based on their function, anatomical location or by the expression of a small number of marker genes (*1*), yet, these classification systems do not fully overlap.

A novel cell type classification system is recently gaining popularity, and it is mostly based on clustering of global molecular profiles as defined by transcriptomic analysis of single cells. This method can answer central questions in kidney biology and disease pathogenesis. First, unbiased single cell clustering can redefine kidney cell types based only on their global transcriptome patterns. Thus far, such analysis has been applied to immune system (*2*), brain (*3*), retina (*4*) cells and even whole multicellular organisms such as drosophila embryos (*5*) and C. elegans (*6*). These experiments have identified novel cell types as well as catalogued marker genes for previously defined cell types, indicating that this approach has the power to redefine kidney cell types.

Second, it is necessary to exploit novel molecular genetic techniques to pinpoint the molecular mechanisms known to cause common kidney diseases (*7*, *8*). This is because kidney pathologies are lumped together by their temporal patterns (acute or chronic) or by their target structure (glomerular vs tubular), which obscures the underlying kidney biology and prevents targeted precision medicine. Further, current pathological measurements cannot distinguish primary cell autonomous responses from secondary cell non-autonomous responses. Previously obtained bulk transcriptome profiles have only been able to generate read-outs for the predominant cell population such as proximal tubule (*9*). In contrast, single cell analysis can determine cell-type specific changes.

Third, single cell analysis may be able to identify fluctuating states of the same cell type. It is generally believed that terminally differentiated cells have limited plasticity and most cell plasticity has been observed in the context of progenitor cell differentiation in the adult state (*10*). In kidney, cell type transition has been only studied in development and between subtypes of intercalated cells (*11*–*14*).

Fourth current models of kidney disease cannot distinguish primary cell autonomous responses from secondary cell non-autonomous responses; single cell specific gene expression profiles could identify the cellular readout of disease associated gene mutations. In sum, using single cell genomic approaches, we have tied genomic discovery with cellular discoveries and molecular and cellular understanding of chronic kidney disease.

### Single cell profiling and unbiased clustering of mouse kidney cells

First, we catalogued mouse kidney cell types in an unbiased manner using droplet-based singlecell RNA sequencing (*15*). We isolated and sequenced 57,979 cells from whole kidney cell suspensions of 7 healthy male mice. Using stringent quality controls (*15*), we further analyzed 43,745 cells. Dimension reduction identified 16 distinct cell clusters consisting of 24 cells to 26,482 cells in each cluster (the clusters were censored for minimum of 20 cells) (**Fig. 1A**).

We have performed several important quality control analyses to validate our results. First we made sure that cells from the 7 different batches (or mice) were distributed evenly in all 16 clusters and each cluster contained cells from more than 4 experiments (**fig. S1**). Next, we separated cells based on the percent of mitochondrial reads (**fig. S2A**). We found that the overall mitochondrial gene percentage slightly influenced the clustering when all cells were analyzed together. On the other hand cells with high or low mitochondrial content showed similar cell clustering patterns, when clustered separately, indicating that the increased mitochondrial gene count is inherent to the specific (proximal and distal tubule) cell types in the kidney (**fig. S2, B and C**). By testing different clustering methods we found that most methods identify similar cell groups (**fig. S3**). It seems that the cells expressing same marker genes were clustered together but some of methods showed less clear separation. Finally, we show that decreasing the cell number from 40,000 to 10,000, 3,000 and 1,000 cells (**fig. S4A**) was associated with increased uncertainty in cell cluster identification and loss of rarer cell types (**fig. S4, B and C**).

**Fig. 1.**
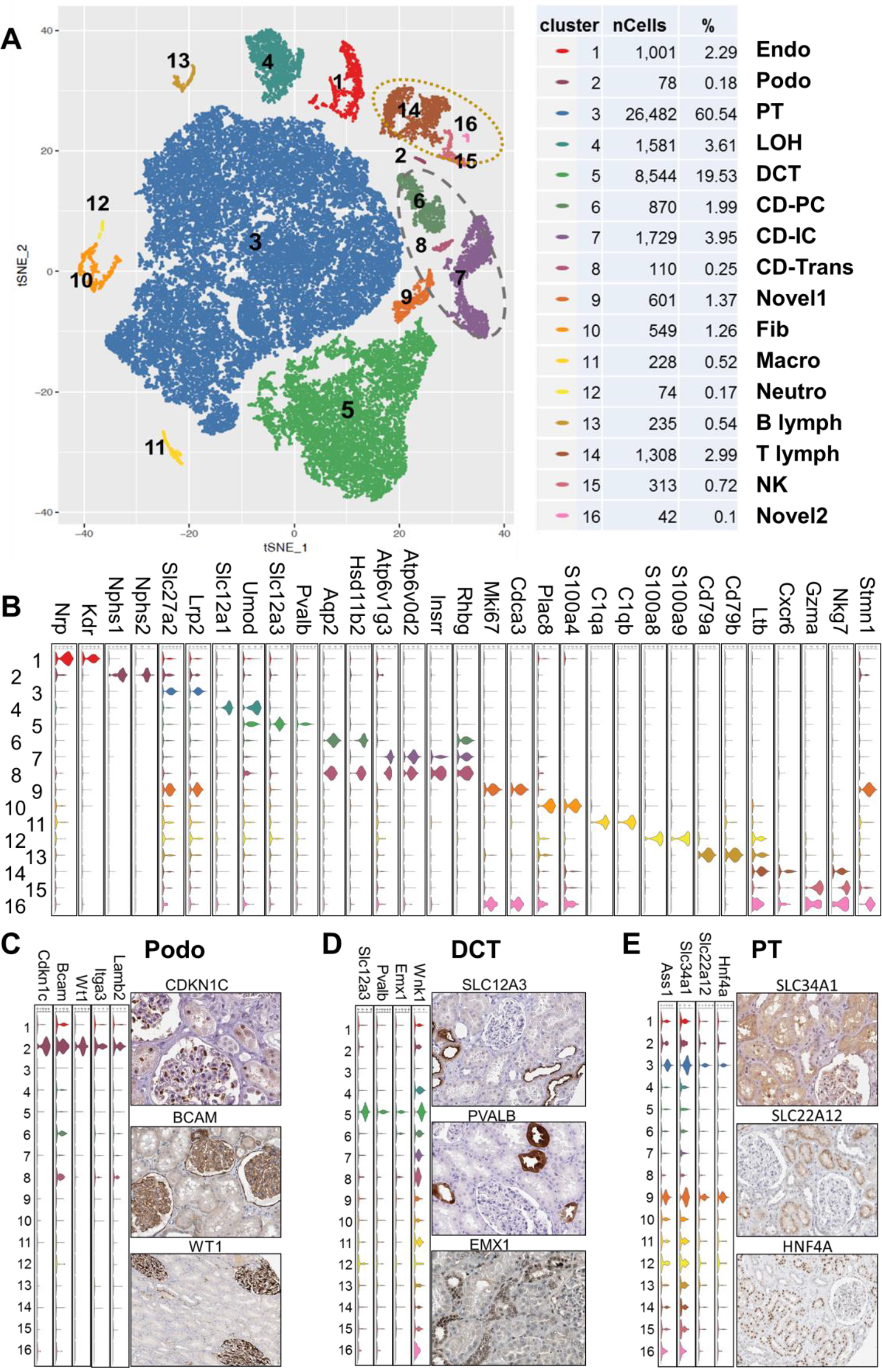
Cell diversity in mouse kidney delineated by single cell transcriptomic analysis. (**A**) tSNE map showing 16 distinct cell types identified by unsupervised clustering. Color code, number of cells, % of cells and assigned cell types are summarized in the right panel. Endo: endothelial, Podo: podocyte, PT: proximal tubule, LOH: loop of henle, DCT: distal convoluted tubule, CD-PC: collecting duct principal cell, CD-IC: CD intercalated cell, CD-Trans: CD transient cell, Fib: fibroblast, Macro: macrophage, Neutro: neutrophil, NK: natural killer cell. (**B**) Violin plots showing the expression level of representative marker genes across the 16 clusters. Y-axis is log scale normalized read count. (**C-E**) Left panel is the violin plot showing the expression level of marker genes for (**C**) Podo, (**D**) DCT and (**E**) PT across the 16 clusters. Y-axis is log scale normalized read count. Right panel is the immunohistochemistry (IHC) data for kidney tissue from The Human Protein Atlas.

### Classification of kidney cells based on cell-type specific marker genes

To define the identity of each cell cluster, we generated cluster-specific marker genes by performing differential gene expression analysis (**Fig. 1B**, **fig. S5 and table S1**) (*15*). In many cases, the unbiased cell cluster identifier marker genes were already known cell-type specific markers, such as the vascular endothelial growth factor receptor 2 (Kdr) for endothelial cells, Nephrin (Nphs1) and Podocin (Nphs2) for podocytes, Na/K/2Cl cotransporter (Slc12a1) for loop of Henle and thiazide sensitive sodium-chloride cotransporter (Slc12a3) for distal convoluted tubule (**Fig 1B**). Immune cells and endothelial cell clusters separated from epithelial cells. The ureteric bud-(cluster 6-8) and metanephric mesenchyme-derived epithelial clusters (cluster 2-5) formed closer clusters (**Fig. 1A**). While some of the markers have already been known, we identified a large number of novel markers, including Cdkn1c and Bcam for podocytes (**Fig. 1B, 1C and table S1**). The cell-type specific protein expressions of such markers were supported by immunohistochemistry analysis (**Fig. 1C-E**).

To reliably assign cell types to cell clusters, we performed careful validation of our results. First, we correlated our gene expression results with bulk RNA sequencing data of microdissected rat kidney segments (**fig. S6**) and microarray data of human immune cell types (**fig. S7**). This comparison highlighted the key advantages of the single cell approaches compared to the analysis of single structure, because single cell analysis demonstrates that glomeruli (endothelial and podocyte) and collecting duct (principal and intercalated cells) consisted of multiple cell types whereas single structure analysis provides only aggregated transcriptomic data from individual cell types. To further validate the clustering analysis results, we used Nphs2^Cre^mTmG, Scl^Cre^mTmG and Cdh16^Cre^mTmG mice as reporter lines to mark podocytes, endothelium and tubule cells. The GFP expression in these models confirmed the proposed cell identity of our cell clusters (**fig. S8**). Altogether, our single cell transcriptome atlas provides a molecular definition for 13 previously defined kidney or immune cell types and 3 novel cell types.

### Mendelian disease genes show cell-type specificity

Next, we tested the hypothesis that genetic kidney diseases that show similar phenotypic changes originate from a single cell type conversely cell function can be inferred by mapping loss of function human gene mutations to these single cell types. We found that 21 out of 29 monogenic gene mutations associated with proteinuria development were expressed in only one cell type of the kidney; the podocyte (**Fig. 2A** **and fig. S9**). Even though earlier studies have implicated the key role of other cell types such as endothelial cells and proximal tubules in proteinuria development, and changes in these cells types can be seen in patients with such conditions, our results unequivocally show the key contribution of podocytes to proteinuria. We found that genes associated with renal tubule acidosis were only expressed in the intercalated cells (IC) of the collecting duct, confirming the major role of intercalated cells in acid secretion (**Fig. 2A**). Genes that are associated with the Mendelian-form of blood pressure (BP) regulation such as Slc12a1, Wnk1, Wnk4 and Klhl3 were also expressed in the collecting duct in addition to the distal convoluted tubule (**Fig. 2A and fig. S9**).

Following the same logic we mapped putative complex trait disease genes such as blood pressure (BP), chronic kidney disease (CKD) and metabolite traits (**Fig. 2B** and **fig. S10**). Surprisingly, most genes implicated in these traits were expressed only in a single cell type. Indeed, very few genes showed a broader multiple cell type expression. Genes associated with plasma metabolite levels and CKD showed strong enrichment for proximal tubule specific expression, while blood pressure associated genes were also enriched in the collecting duct cells. In summary our single cell transcriptome maps can provide valuable insight into the expression of genetic variants, to highlight the responsible cells, and to assign cellular functions.

**Fig. 2.**
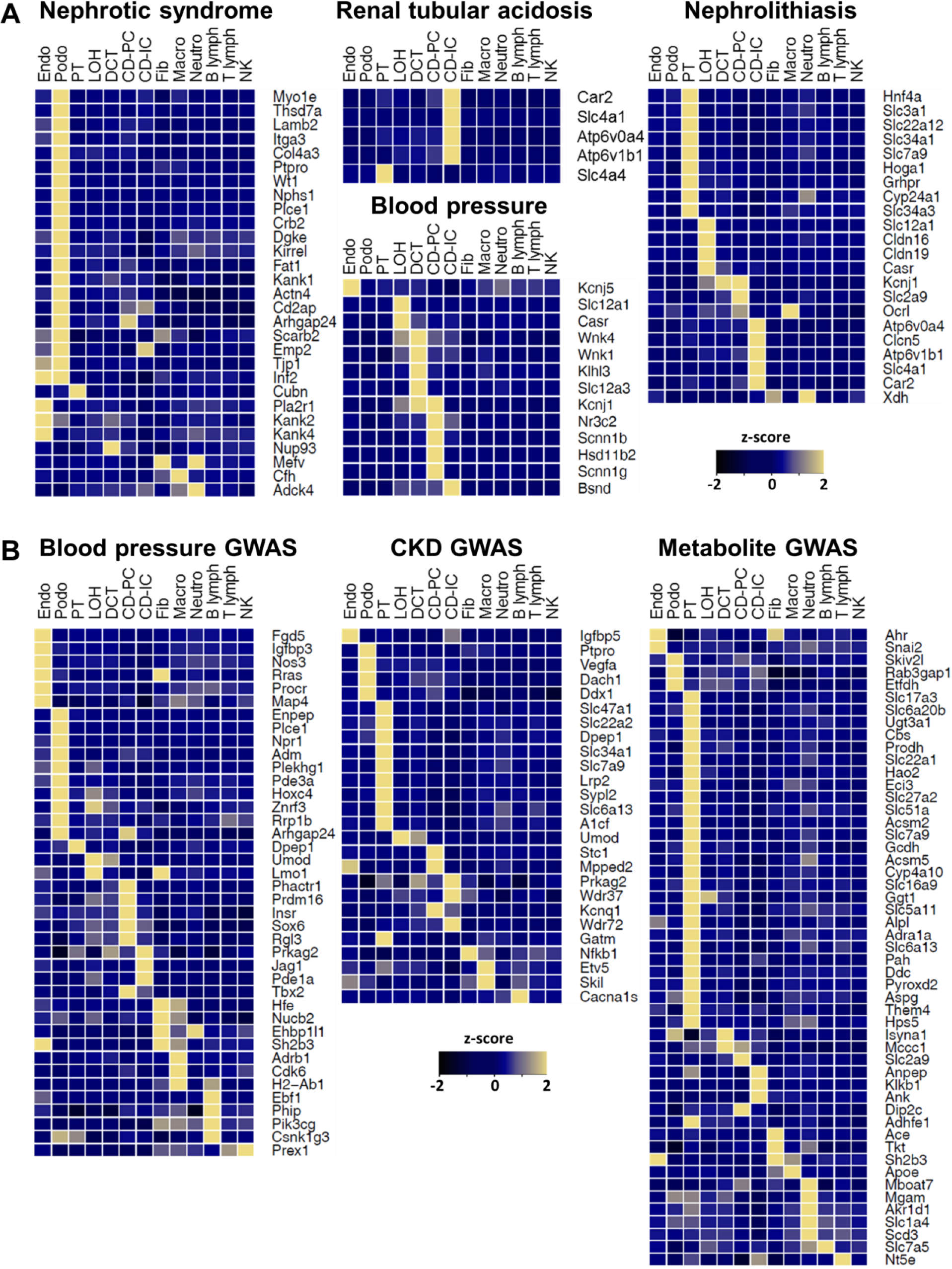
Molecular dissection is applied to map the disease-associated genes. (**A and B**) Average expression level of the human (**A**) monogenic disease genes and (**B**) GWAS genes. Mean expression values of the genes were calculated in each cluster. Color scheme is based on z-score distribution. The cell types only assigned to known kidney cells and immune cells, and the only genes whose maximum z-scores are more than 2 are shown in the heatmaps. Full list of cell types and genes are shown in fig. S, 9 and 10.

### Identification of a novel cell type in the renal collecting duct

The kidney collecting duct differs from all other kidney epithelia since it originates from the ureteric bud and not the metanephric mesenchyme. This compartment is known to be comprised of at least two molecularly and functionally distinct cell types: the principal cells (PC), responsible for sodium, water reabsorption and potassium secretion, and the intercalated cells (IC) responsible for acid secretion. Our analysis identified aquaporin 2 (Aqp2) and H+-ATPase (Atp6v1g3) as the key marker genes for clusters 6 and 7, defining these clusters as PC and IC cells (**Fig. 3A** **and table S1**). Immunohistochemistry analysis confirmed the exclusive expression of these markers in collecting duct cells (**Fig. 3A**). The intercalated cell cluster separated into two subclusters and anion exchanger 1 (Slc4a1) and pendrin (Slc26a4) were key marker genes for these subclusters (**Fig. 3A**). These results are consistent with prior results indicating that intercalated cells can be subdivided to typeA and typeB IC cells based on Slc4a1 and Slc26a4 expressions.

Unexpectedly, our single cell profiling identified another cell cluster. This cell cluster (cluster 8) expressed markers of both IC and PC cells (**Fig. 3**, **A and B**) in addition to cell-type specific markers such as Car12, Rhbg, Insrr and Clcnkb (**Fig. 3**, **B and C**). To confirm the existence of this cell type, we performed double immunofluorescence staining with Aqp2 and Atp6b1 (**Fig. 3D**).

To further understand the identity of this novel cell type, we utilized the Monocle analysis toolkit to perform cell trajectory analysis using pseudotime reconstitution of clusters 6-8 (*15*). We found that the novel cells were located between PCs and ICs suggesting that cluster 8 is a transitional cell type but not a progenitor which is expected to be located in beginning of the either branches (**Fig. 3E**). Furthermore, cell trajectory analysis clearly separated IC and PC cells into their subtypes as previously identified (*11*, *16*) verifying its performance (**fig. S11**). These results indicate that the collecting duct contains not only PC and IC cells but a third distinct transitional cell type, raising the possibility that IC and PC cells represent two ends of a spectrum of cellular phenotypes and that they interconvert.

### Fluorescent lineage tracing confirms collecting duct cell plasticity

We next examined whether our aforementioned computational identification of transitional cells was visible by conventional *in vivo* lineage tracing. We generated mice that carry a lineage tag in differentiated principal cells (Aqp2^Cre^mT/mG) and in differentiated intercalated cells (Atp6^Cre^mT/mG) (**Fig. 3**, **F and G**). We performed triple immunofluorescence labeling in these animals by staining for GFP (all cells of a specific marker origin), Aqp2 (principal cells) and Atp6ase (intercalated cells). As expected, we found that most of the GFP-positive cells were also Aqp2-positive in the Aqp2^Cre^mT/mG mice, a subset of the GFP positive cells only expressed Atp6b1, an IC marker, but not Aqp2 and smaller subset was double positive for Atp6b1and Aqp2 (**Fig. 3F**). Among the GFP-positive cells, 61.6% of the cells were Aqp2-positive, 29.2% were Atp6b1-positive and only 9.2% of the cells were positive for both (**Fig. 3F**). Similar analysis performed in the Atp6^Cre^mT/mG mice showed that Aqp2-positive and Atp6b1-positive (transitional cells) or Atp6b1-negative true PC cells can originate from Atp6b1-positive IC cells (**Fig. 3G**). To determine if cell proliferation could be responsible for this cell plasticity, we calculated the expression levels of cell-cycle regulated genes in the single cell transcriptome map. We found that only cluster 9 and 16 (novel cell types 1 and 2), but not any of the collecting duct clusters, showed higher expression levels of the cell-cycle genes (**fig. S12**). This suggests that cluster 8 is likely to be a transitional cell population and not a proliferating progenitor cell type. In summary, *in vivo* lineage tagging analysis confirmed an interconversion of PC and IC cells in the collecting duct via a novel transitional cell type.

**Fig. 3.**
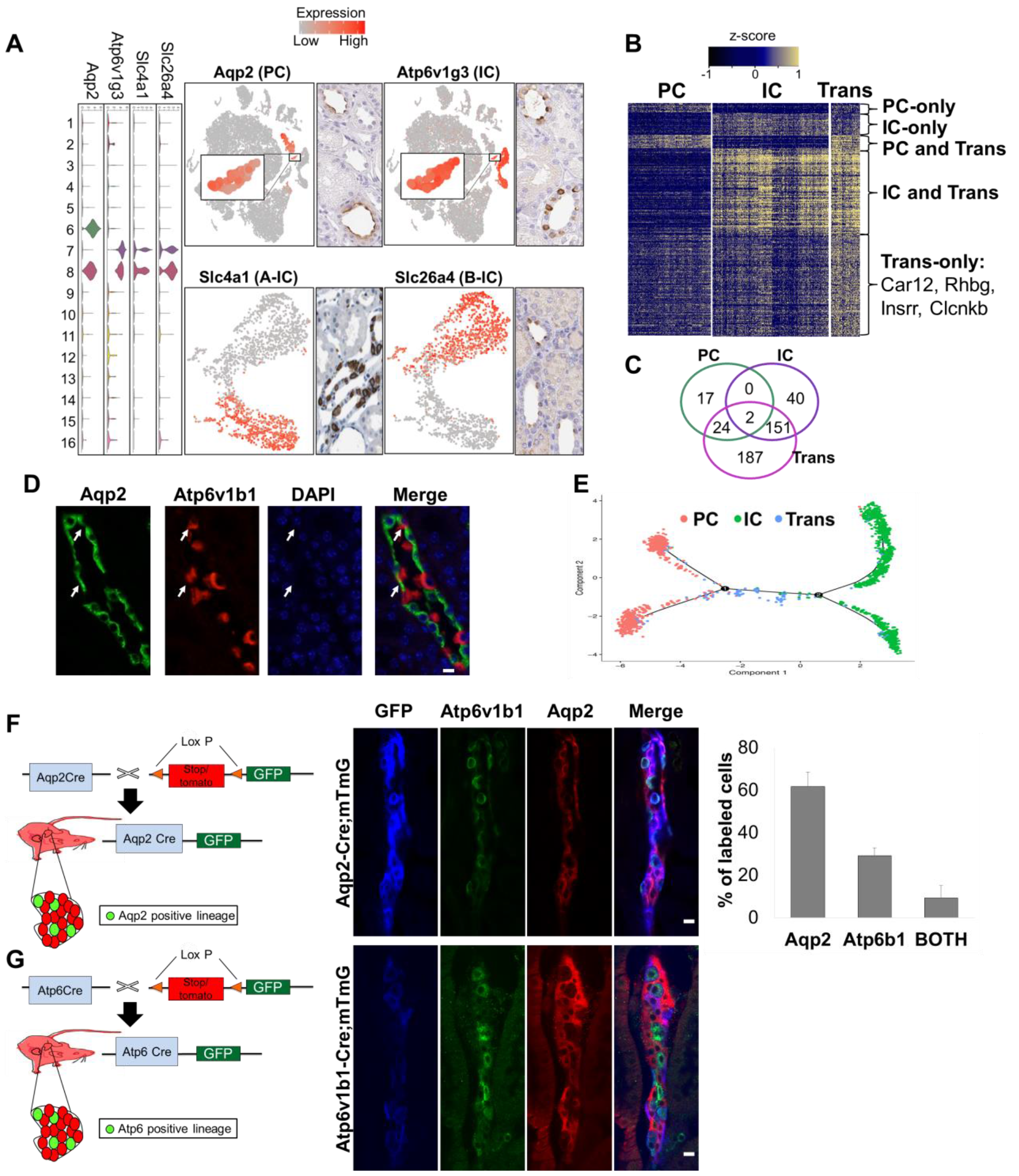
Identification of a novel transitional cell type in the collecting duct. (**A**) Left panel is the violin plot showing the expression level of marker genes across the 16 clusters. Y-axis is log scale normalized read count. Right panel is the tSNE plot showing gene expression level, and the immunohistochemistry (IHC) data from The Human Protein Atlas showing protein expression patterns. (**B**) Heatmap showing the expression level of differentially expressed genes in collecting duct cell types compared to all 16 cell clusters. Color scheme is based on z-score distribution. Example genes for trans-only differentially expressed genes are listed. (**C**) Venn diagram showing the overlaps of DEGs between PC, IC and the novel cell type. (**D**) Immunofluorescent staining for Aqp2 (PC marker), Atp6v1b1 (IC marker) and DAPI in the renal collecting duct. Scale bar: 10um. (**E**) Ordering single cells along a cell conversion trajectory using Monocle. Three collecting duct cell clusters were used for ordering and plotted on low dimensional space with different colors. (**F**) Immunofluorescent staining for GFP, Atp6v1b1 and Aqp2 in mouse model used for lineage tracing of Aqp2-positive cells. Scale bar: 10um. Right panel is the quantification result of labeled cells among GFP positive cells. n=3. (**G**) Immunofluorescent staining for GFP, Atp6v1b1 and Aqp2 in mouse model used for lineage tracing of Atp6-positive cells. Scale bars: 10um

### Collecting duct cell plasticity is mediated by Notch signaling and is likely important in disease development

For further analysis of collecting duct cell plasticity, we identified genes of which the expressions change as the cell transition progresses (**fig. S13, A and B**) (*15*). Genes with increased levels in PCs were associated with cell adhesion, water homeostasis and salt transport while the genes upregulated in ICs were associated with ATP hydrolysis/synthesis coupled proton transport and oxidation-reduction processes (**fig. S13C**), highly consistent with these cell functions. In addition, one of the key signaling pathways that showed differential expression during the cell transition of the collecting duct cells was Notch signaling. Notch ligands such as Dll4 and Jag1 were highly expressed in ICs and their expression was low in PC cell (**Fig. 4A**). In contrast, PCs highly expressed Notch2, Notch3 receptors, and their target genes Heyl and Hes1, suggesting that PCs are the Notch signal receiving cells in collecting duct. Immunoflourescence studies confirmed the exclusive ligand (Jag1) expression pattern in the IC cells (**Fig. 4B**).

Next, we examined whether Notch signaling drives the IC/PC cell transition. We generated the Pax8rtTA/TRENICD mice, which enables inducible transgenic expression of the Notch receptor (intracellular domain) specifically in adult (differentiated) renal tubule cells (**Fig. 4C**). We found that the number of cells expressing the PC cell marker Aqp2 was increased, while the number of cells expressing the IC cell marker Atp6b1 was reduced in this mouse model (**Fig. 4C**). We also performed in silico cell type deconvolution of bulk RNA profiling data to estimate cell type fractions in control and Pax8rtTA/TRENICD mice (*15*). This analysis confirmed the increase in PC over IC cell ratio in Pax8rtTA/TRENICD mice (**Fig. 4D**). Collectively, our data indicate that Notch receptor expression and signaling is sufficient to drive the PC cell fate even in the adult collecting duct.

Because increased Notch receptor expression have been reported in patients and animal models with kidney disease (*17*), we examined whether disease states disrupt IC/PC balance and cell transition. In kidneys of mice with folic acid induced (FA) kidney disease, we found an increase in PC cells and a decrease in IC cells when compared to control mice (**Fig. 4E**). Computational cell deconvolution analysis of bulk RNA sequencing data obtained from control FA kidney disease model supported the shift toward PC cell fate (**Fig. 4F**). Using cell markers identified in mice, we also performed computational deconvolution analysis of 91 human patient samples. Again, we found that PC fraction increased in diseased samples while IC cell fraction decreased (**Fig. 4G**), consistent with increased Notch signaling and HES1 expression.

Finally, we analyzed whether the increased IC to PC transition in CKD is associated with functional consequences. IC cells are uniquely associated with acid secretion in the kidney and genes that cause metabolic acidosis are exclusively expressed in this cell type (**fig. S13C and Fig. 2A**). We found that total CO_2_ levels (composite measure of serum bicarbonate and pCO_2_) were significantly reduced in the FA induced mouse kidney disease model, consistent with metabolic acidosis (**Fig. 4H**). In summary, IC to PC transition appears to be mediated by Notch ligand and receptor expression and shifted toward the PC cell fate, and this correlates with metabolic acidosis in kidney disease.

**Fig. 4.**
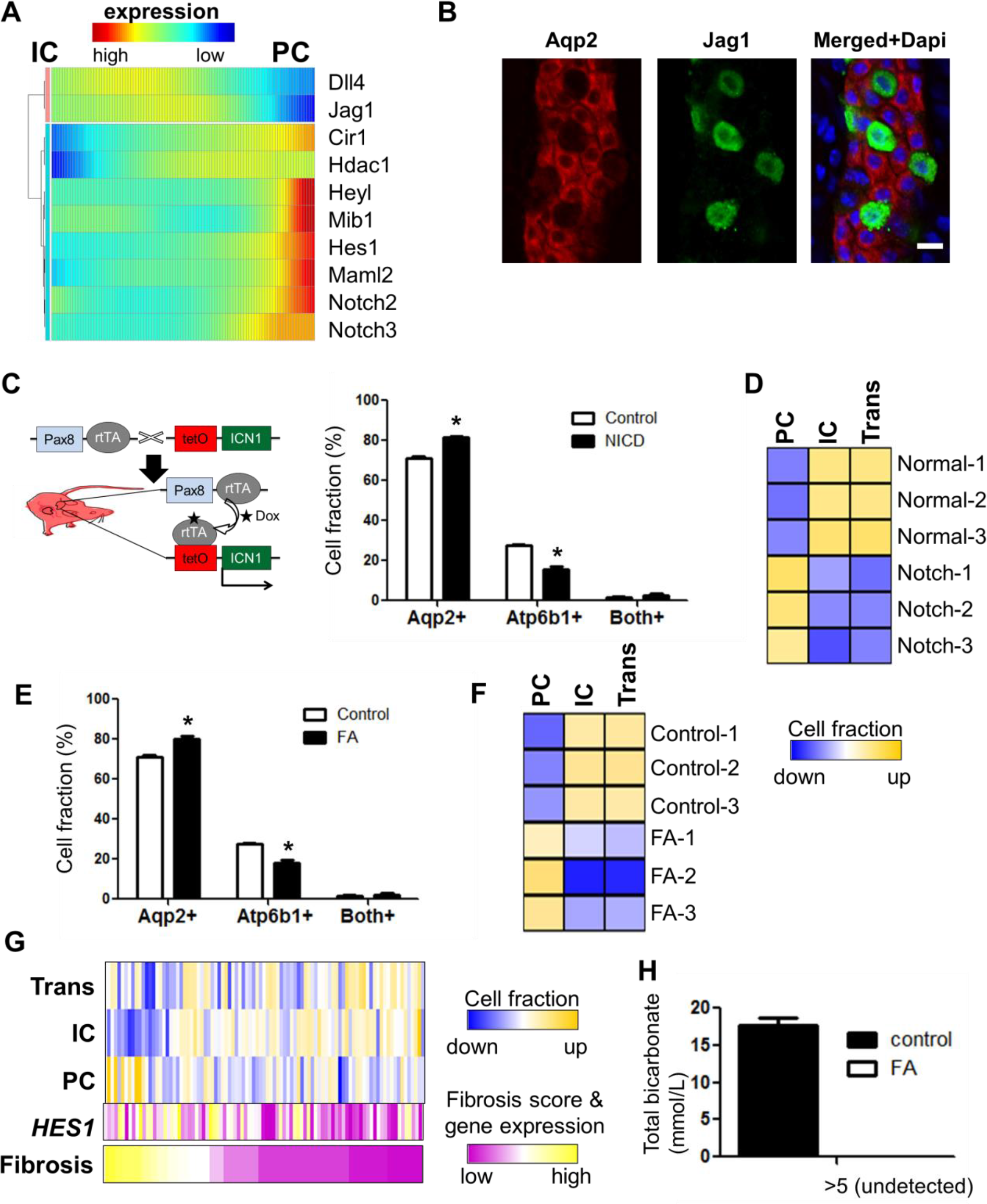
IC to PC transition is mediated by Notch signaling and increased in disease condition. (**A**) Transcriptional dynamics of Notch signaling genes during cell conversion. Cells are ordered in pseudotime and color represents expression levels. (**B**) Immunofluorescent staining for Aqp2 and Jag1 in the renal collecting duct (**C**) Mouse model used to activate Notch signaling in adult kidney tubules. Right panel is the immunofluorescent quantification result of labeled cells with Aqp2 and Atp6b1 in control and Notch Tg mice. n=3. Asterisk represents significant difference; p-value < 0.01. (**D**) Deconvolution of mouse kidney bulk RNA profiling data. Wild type (WT) and intracellular domain of the Notch (NICD) protein overexpression samples were used for analysis. (**E**) Immunofluorescent quantification result of labeled cells with Aqp2 and Atp6b1 in control and FA mice. n=3. Asterisk represents significant difference; *p*-value < 0.01. (**F**) Deconvolution of mouse kidney bulk RNA profiling data. Control and kidney samples from FA injected mice were used for analysis. (**G**) Deconvolution of human kidney microarray data. Human kidney samples from normal and CKD patients (n=91) were used for analysis. The histological fibrosis scores (from 0 to 100) and *HES1* expression levels for the corresponding samples are also shown. (**H**) Total bicarbonate level in blood from control and FA mice. n=5.

## Discussion

Cells are the fundamental units of a multicellular organism. Description of kidney cell types is more than 100 years old and dates back to the invention of the microscope. Since then, a functional kidney cell annotation has been developed that is mostly based on the transport function of the kidney. Here, we provide a new molecular definition of kidney cell types by sequencing gene expression in 57,979 cells. At this resolution, we distinguished 16 major cell types and defined these cells by quantitative gene expression. By performing comprehensive comparative analysis we were able to identify almost all previously described cell types in addition to 3 novel cell types. While a few prior studies have attempted single cell sequencing analysis of kidney cells they were limited by the low cell count of patient samples (*18*), sorted segments or fetal mouse kidney (*19*), and thus were unable to generate a comprehensive assessment.

This kidney cell atlas provides a remarkable insight into kidney function and disease pathogenesis. It demonstrates that genetic diseases of the kidney are referable to a single cell types. While prior transcriptomic studies on kidney disease samples identified changes in multiple cell types, our data indicates that loss of function gene mutations localize to single cell types and disease-causing genes show exquisite cell-type specificity. Our analysis of complex trait genes indicates a similar pattern of single cell-type specific expression of disease-associated genes. It seems that each cell group has a unique (non-redundant) function in the kidney and therefore equipped with a unique expression profiles. Genetic analysis can therefore help to identify disease causing cell types and cell-type specific function.

Our data also highlights the previously underappreciated role of the kidney collecting duct system in health and disease states. Genes with mutations associated with acidosis, CKD and BP are specifically localized to this kidney segment. Furthermore, one of the most striking results of our single cell analysis was the identification of a novel distinct cell type in this kidney segment relating to the well-known IC and PC cells. Computational and lineage tagging analyses indicate that these are most likely transitional cell types. The PC cells are irreplaceable as they are responsible for sodium and water balance, and are involved in the final regulation of serum potassium levels (*16*). Elevated serum potassium results in cardiac conduction abnormalities and the cause of death in people with kidney failure. On the other hand, by regulating the respiratory rate the body can partially compensate for acid accumulation with little acute disruption of serum pH. Perhaps this rationale explains PC cell fate preservation compared to IC cell preservation in disease states.

To conclude, we have generated a comprehensive cell atlas for the kidney, identified cell-type specific markers along with novel cell types, and have uncovered unexpected cell plasticity. The information gained from this study will be essential for further understanding of kidney homeostasis and disease development.

